# GFP fusion protein with embedded foreign peptide

**DOI:** 10.1101/855908

**Authors:** K.A. Glukhova, V.G. Klyashtorny, B.S. Melnik

**Affiliations:** Institute of Protein research RAS, Institutskaya, 4, Pushchino, 142290, Russian Federation

## Abstract

From the point of view structural biology and protein engineering the green fluorescent protein (GFP) is an exceptionally attracting object. The tertiary structure of GFP is quite unique: it reminds a “cylinder” or a “barrel” consisting of beta-layers that contains an alpha-helix inside. The “barrel” is a special container for an alpha-helix serving to protect the latter from the influence of the surroundings. Therefore a reasonable question arises whether the “barrel” can function as a container for preservation and isolation of other peptides. The alpha-helix itself contains hydrophilic amino acids, whereas inside the barrel there are many molecules of bound water. We supposed that the central alpha-helix of green fluorescent protein could be substituted for foreign peptide. In this study we checked the possibility for creation of such a system on base of GFP, where the toxic peptide is isolated from the environment inside the protein. The modification of green fluorescent protein was carried out. An antimicrobial peptide was inserted into the central alpha-helix. The results of our experiments show that such a chimeric protein is compact, soluble and non-toxic for the producing cell culture, but its structure is destabilized. The obtained data show that the idea of use of green fluorescent proteins as a «container» for storing foreign peptides could be realized.

## Introduction

Antimicrobial peptides (AMPs) comprise amphipathic molecules consisting mainly of positively charged and hydrophobic amino acids. AMPs interact with negatively charged phospholipids of bacterial plasma membrane and form pores in the membrane, which leads to the loss of membrane potential resulting in cell death. The peptides affect a wide range of pathogens, such as bacteria, fungi, viruses [1]. Studying AMPs and their therapeutical implementation is limited by a number of problems related to their production and application. For instance, antibacterial activity impedes their production in bacterial culture. Alternative production pathways, such as eukaryotic producers, chemical synthesis or isolation from natural sources, increase the production costs significantly. Modern approaches are based on decreasing AMP toxicity via their synthesis in inactive form. In this way, the peptides are produced in bacterial culture in form of chimeric proteins where auxiliary proteins or peptides (thioredoxin, SUMO, PurF, etc.) neutralize the toxicity of AMPs [2].

Another problem is related to fast degradation of the peptides after their injection into the organism. To address this problem, variants of peptide protection via their encapsulation into liposomes and polymeric particles are proposed [3].

The optimal solution for both problems could be use of an AMP-carrier resistant to proteases and capable of complete peptide isolation from the environment during its synthesis and targeting to the site of interest in the body. From this point of view, green fluorescent protein (GFP) is one of the promising objects. GFP consists of 238 amino acids folded into 6 alpha helices and 11 beta barrels linked by loops. The antiparallel beta sheets form a cylinder where an alpha-helix with a chromophore is placed. The beta barrel is supposed to be a protector of the chromophore from fluorescence quenchers and proteases. GFP is known to be resistant to proteases [4]. Despite its tight packing, free space exists inside the protein, where water molecules can be detected [5]. Also, salt ions can be retained inside GFP [6]. The microenvironment of the chromophore is rich in charged amino acid residues, some of them responsible for ion retention inside the barrel.

Such structural features of GFP let us suppose that the central alpha-helix of this protein could be substituted for foreign peptide. Kent et al. have already tried to substitute the central alpha-helix of this protein for xenogenic alpha-helical peptide. The authors had successfully «replaced» the helices of two different proteins. Meanwhile, the proteins remained structured, and fluorescence of mature chromophores was observed [7]. These results confirm our supposal on the possibility of the replacement of the central alpha-helix of GFP.

For alpha helix substitution, we chose the peptide named bactenecin. It is a member of antimicrobial peptides group; it is synthesized by neutrophils and has an activity against Gram-negative bacteria, such as *Escherichia coli, Pseudomonas aeruginosa, Salmonella typhimurium* [8]. The selection was based on the length and position of hydrophilic and hydrophobic amino acids in bactenecin amino acid sequence.

Creation of a complete system for synthesis and targeting of antimicrobial peptides suitable for practical application is a labour-consuming multi-stage task. The goal of the current work was to check the possibility for creation of such a system on base of GFP, where the synthesized toxic peptide is isolated from the environment inside the protein.

## Materials and methods

**Hybrid protein design** was carried out in RasMol program (http://www.rasmol.org). Optimization of hybrid protein was carried out by molecular dynamics method in Gromacs 4.5.3 program [9], CHARMM27 force field. Spatial structure of 1B9C [10] from Protein data bank (PDB: https://www.rcsb.org/) was used as a starting model for molecular dynamics. The model was placed into 65 × 65 × 65 Å orthorhombic water box filled with TIP3P water molecules. The water box was built so that the distance from protein surface in each direction to the edge of water box was at least 12Å. To test the quality of the obtained structures, deviation of stereo-chemical parameters after 270 ns molecular dynamics was calculated. No significant deviation of protein coordinates from starting conformation was observed. Structure of GFP fluctuated around its equilibrium position. Root-mean square deviations for all the atoms of the system did not exceed 3 Å (2 Å for Cα atoms of proteins) in all the trajectories. To analyze the effect of mutations on protein structure, 300 ns molecular dynamics trajectories were simulated at constant temperature, 300K and pressure, 1 atm. All the simulations were carried out using computational resources of Joint SuperComputer Center of the Russian Academy of Sciences (JSCC RAS, www.jscc.ru). The coordinates of the system were saved and analyzed every 1 ps.

### DNA synthesis and protein isolation

DNA fragment encoding the hybrid protein GFP-bactenicin was synthesized by PCR with overlapping primers. cDNA fragment was cloned in expression vector pET28a by NdeI and EcoRI restriction sites. Site-directed mutagenesis was carried out as described earlier [6, 11].

Recombinant proteins were produced in cell cultures of *E.coli* BL21(DE3). Protein synthesis was induced by addition of isopropyl-β-D-1-thiogalactoside (IPTG) to the final concentration of 0.1 mM at OD_600_ = 0.5. The cells were cultivated for 16 h at 18 °C. Cell pellet was resuspended in 20mM Tris-HCl buffer, pH 8.0, supplied with 1mM EDTA, and disrupted by an ultrasonic disintegrator Branson Sonifier. Cell lysates were centrifuged at 20 000 g for 30 min. Supernatant and pellet were analyzed by PAAG electrophoresis in denaturing conditions.

## Results and discussion

A hybrid protein, GFP-bactenecin, with a part of inner alpha-helical GFP moiety being substituted for bactenecin amino acids, was constructed in the work. Three amino acids, S, Y, and G, which form a chromophore in native GFP, were added into bactenecin moiety. Figure 1 shows the structure of GFP and amino acid sequence of GFP alpha-helix, hybrid GFP-bactenecin protein and bactenecin.

**Figure 1.**
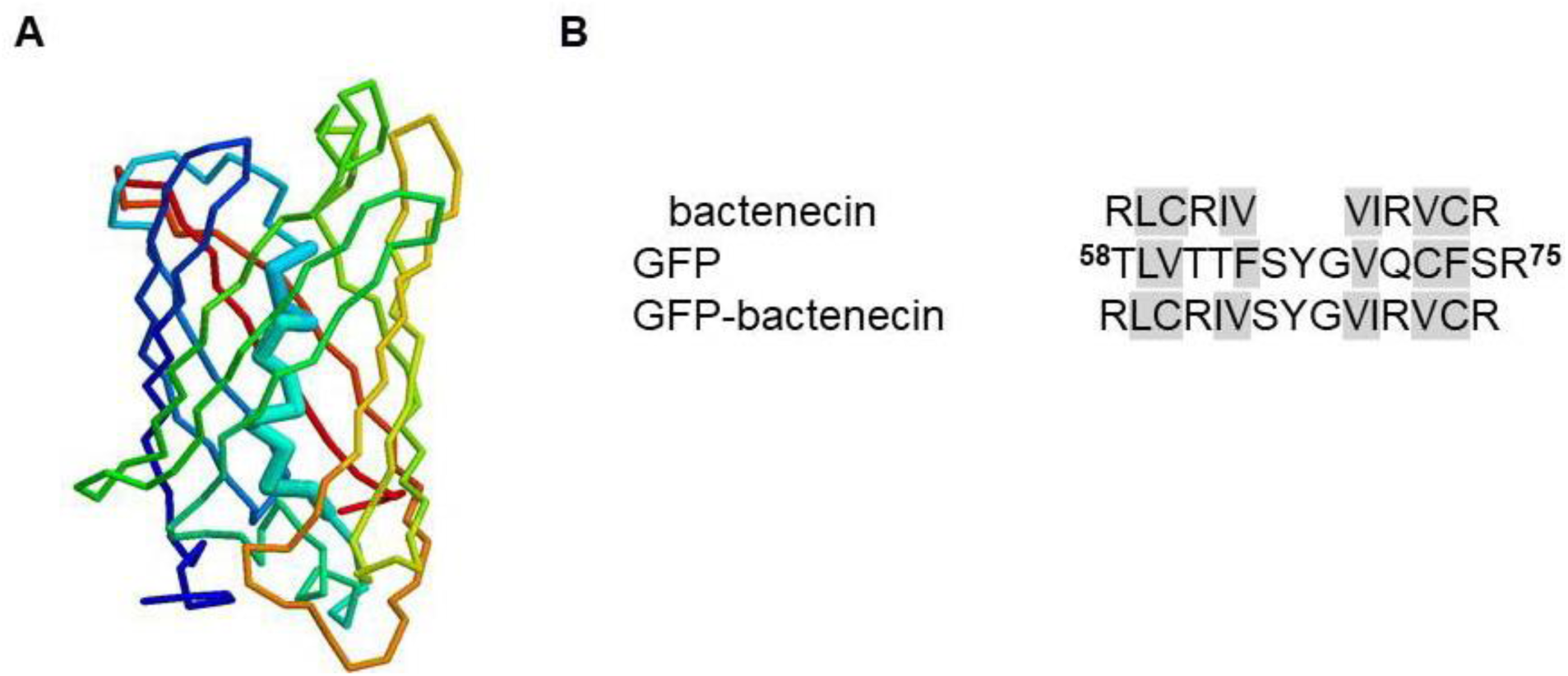
A— 3D structure of GFP; B – sequence alignment of GFP alpha-helix, GFP-bactenecin hybrid protein and bactenecin. Hydrophobic amino acids are highlighted in gray.

Very similar arrangement and alternation of hydrophilic and hydrophobic amino acids specific for alpha-helices of soluble proteins is observed in amino acid sequences of bactenecin and GFP (Fig. 1B). Nevertheless, the designed insertion of bactenecin instead of the central alpha-helix alters the majority of amino acid residues in the center of GFP (Fig. 1A). So, our first task was to reveal the possibility of GFP folding after multiple substitutions in the central alpha-helix.

The hybrid protein was produced in *E.coli* bacterial culture. It was found that GFP-bactenecin is synthesized in a significant amount and it is not toxic for the culture of producer. But the cells had no fluorescence, i.e. the chromophore was not formed properly in the hybrid protein. After ultrasound disintegration of producer cells, GFP-bactenecin was detected in insoluble protein structure (Figure 2). This is the evidence that GFP-bactenecin is unstable and forms aggregates (modification leads to misfolding and aggregation).

**Figure 2.**
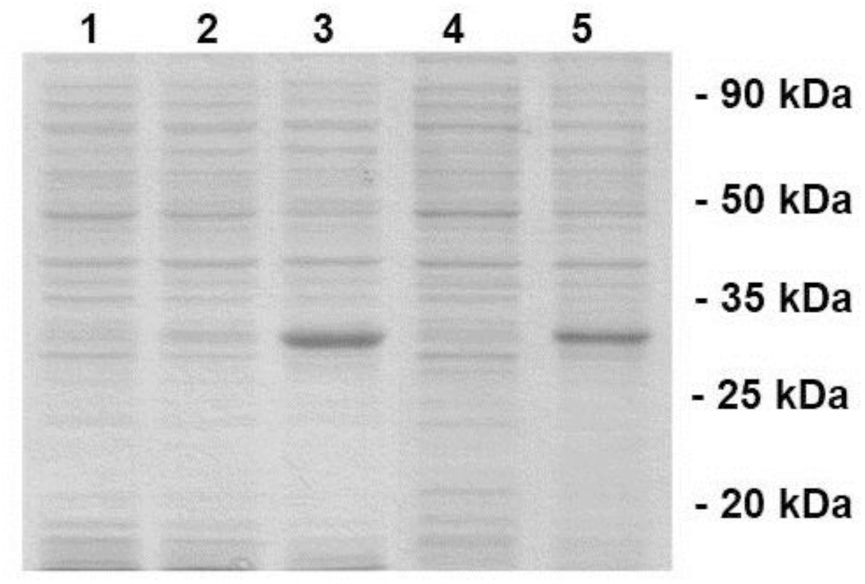
Electrophoregram of cell samples producing GFP-bactenecin. 1 — cells before induction by IPTG; 2 — cells after 2 hours of induction by IPTG; 3 — cells after 4 hours of induction by IPTG; 4 — soluble protein fraction; 5 — insoluble protein fraction.

As wild-type GFP is strongly aggregating at elevated temperature or in presence of denaturants [12; 13], we supposed that the reason of chimeric protein aggregation could be destabilization of the protein structure. To search for mutations stabilizing GFP-bactenecin, improving its folding and thus decreasing aggregation, modeling of GFP structure by molecular dynamics was carried out.

The analysis of GFP-bactenecin structure by molecular dynamics and its comparison with wild-type protein showed that substitutions of three amino acids (F8D, I167D, Q177E) could actually stabilize this protein. We modeled the protein structure with F8D, I167D, Q177E substitutions, and simulation showed that the chosen substitution improve the compactness of the protein and possibly stabilize it.

Figure 3 shows the structure of GFP-bactenecin protein (Figure 3A) simulated by molecular dynamics method and structure of GFP-bactenecin protein with additional stabilizing mutations (Figure 3B). It can be seen that the structure of GFP-bactenecin is «distorted» compared to wild-type protein. Simulation of GFP-bactenecin with additional mutations showed that the chosen mutations could stabilize protein structure.

**Figure 3.**
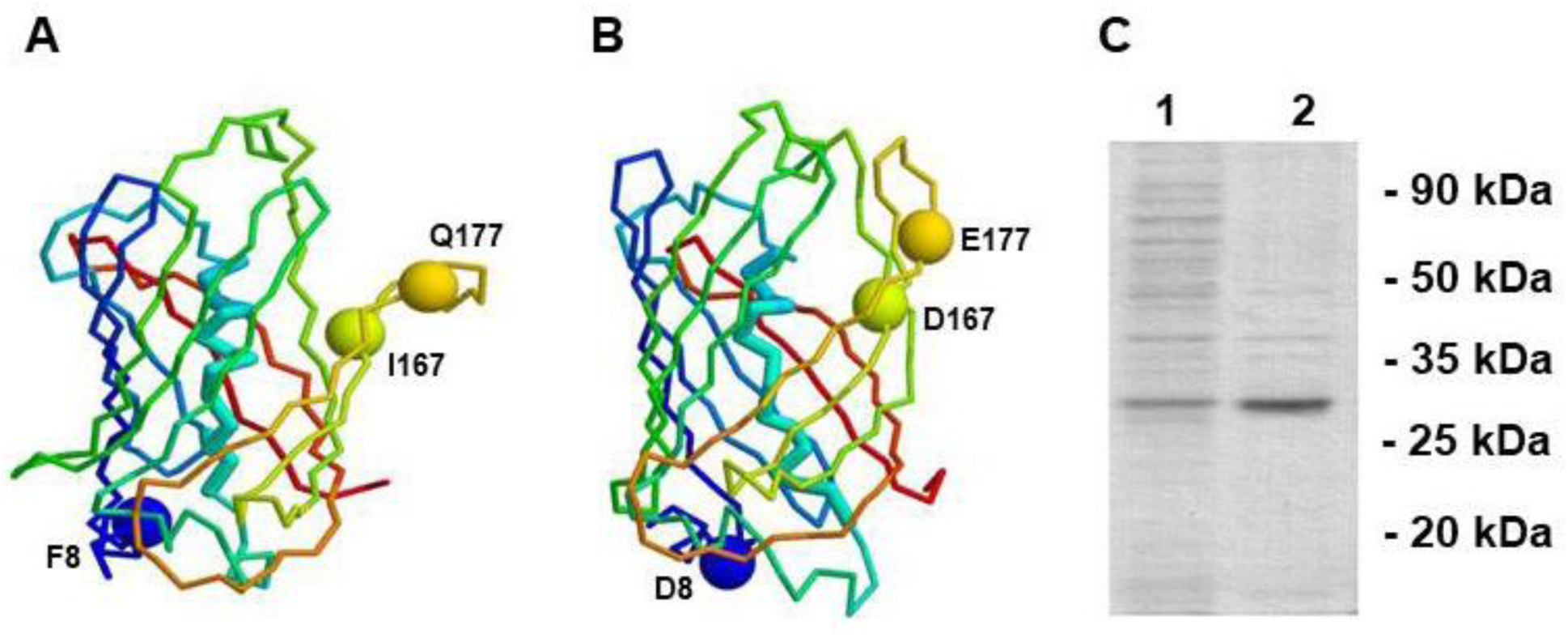
A. Structure of GFP-bactenicin modeled by molecular dynamics. B Optimized structure of GFP-bactenecin protein. C. Electrophoregram of cell samples producing GFP-bactenecin after 4 h of induction by IPTG; 1 — soluble protein fraction, 2 — insoluble protein fraction.

As a result, an optimized GFP-bactenecin variant with stabilizing F8D, I167D, Q177E substitutions was constructed. The optimized protein had no toxic effect on the cells during its production in bacterial culture. Unlike the previous variant, the optimized protein was detected in soluble protein fraction after ultrasound disintegration of the cells (Figure 3C). Thus, the inserted substitutions stabilized the protein structure and reduced aggregation. Unfortunately, neither cell culture, nor isolated protein (from soluble protein fraction) had any detectable fluorescence. This evidences that the chromophore is not formed in the modified GFP.

Nonetheless, the obtained data show that GFP can be used as a «container» for toxic peptides. The studies of this and earlier work [6], as well as theoretical computation by molecular dynamics method, show that the changes in the central alpha-helix and substitution of a large number of hydrophilic or hydrophobic amino acids destabilize the protein, but do not «forbid» this protein to have structure similar to that of wild-type protein.

## Acknowledgements

The authors gratefully thank Anatoly Sergeevich Glukhov for his help during expression vector construction. The work was financially supported by Russian Foundation for Basic Research, grant number 18-34-00313.

## References

1. Hancock R. E., Haney E. F., Gill E. E. The immunology of host defence peptides: beyond antimicrobial activity. Nat Rev Immunol, 2016, vol. 16, pp.321–334. doi: 10.1038/nri.2016.29.

2. Li Y. Carrier proteins for fusion expression of antimicrobial peptides in Escherichia coli Biotechnol. Appl. Biochem, 2009, vol. 54, pp. 1–9. doi: 10.1042/BA20090087.

3. Urbán P., Valle-Delgado J. J., Moles E., Marques J., Díez C., Fernàndez-Busquets X. Nanotools for the delivery of antimicrobial peptides. Curr Drug Targets, 2012, vol. 13, pp. 1158–1172. doi: 10.2174/138945012802002302.

4. Bokman S. H., Ward W. W. Renaturation of Aequorea gree-fluorescent protein. Biochem Biophys Res Commun, 1981, vol. 101, pp. 1372–1380. doi: : 10.1016/0006-291X(81)91599-0

5. Ormö M., Cubitt A. B., Kallio K., Gross L. A., Tsien R. Y., Remington S. J. Crystal structure of the Aequorea victoria green fluorescent protein. Science, 1996, vol. 273, pp. 1392–1395. doi: 10.1126/science.273.5280.1392.

6. Glukhova K. F., Marchenkov V. V., Melnik T. N., Melnik B. S. Isoforms of green fluorescent protein differ from each other in solvent molecules ‘trapped’ inside this protein. J Biomol Struct Dyn, 2017, vol. 35, pp. 1215–1225. doi: 10.1080/07391102.2016.1174737.

7. Kent K. P., Oltrogge L. M., Boxer S. G. Synthetic Control of Green Fluorescent Protein J Am Chem Soc, 2009, vol. 131, pp. 15988–15989. doi: 10.1021/ja906303f.

8. Wu M., Hancock R. E. Interaction of the Cyclic Antimicrobial Cationic Peptide Bactenecin with the Outer and Cytoplasmic Membrane, J Biol Chem, 1999, vol. 274, pp. 29–35. doi: 10.1074/jbc.274.1.29.

9. Abraham M. J., Murtola T., Schulz R., Páll S., Smith J.C., Hess B., Lindahl E. GROMACS: High performance molecular simulations through multi-level parallelism from laptops to supercomputers. SoftwareX, 2015, vol. 1, pp. 19–25. doi: 10.1016/j.softx.2015.06.001.

10. Battistutta R., Negro A., Zanotti, G. Crystal structure and refolding properties of the mutant F99S/M153T/V163A of the green fluorescent protein. Proteins, 2000, vol. 41, pp. 429–437. doi: 10.1002/1097-0134(20001201)41.

11. Melnik T.N., Povarnitsyna T.V., Glukhov A.S., Melnik B.S. Multi-state proteins. Approach allowing experimental determination of the formation order of structure elements in the green fluorescent protein. PloS ONE 2012. 7 (11): e48604. doi: 10.1371/journal.pone.0048604.

12. Tsien R. Y. The green fluorescent protein. Annu Rev Biochem, 1998, vol. 67, pp.509–544. doi: 10.1146/annurev.biochem.67.1.509.

13. Stepanenko O. V., Stepanenko O. V., Kuznetsova I. M., Verkhusha V. V., Turoverov K. K. Beta-Barrel Scaffold of Fluorescent Proteins: Folding, Stability and Role in Chromophore Formation. Int Rev Cell Mol Biol, 2013, vol. 302, pp. 221–278. doi: 10.1016/B978-0-12-407699-0.00004-2.

